# A Novel Y-Shaped, S-O-N-O-S-Bridged Crosslink between Three Residues C22, C44, and K61 Is a Redox Switch of the SARS-CoV-2 Main Protease

**DOI:** 10.1101/2022.04.29.490044

**Authors:** Kai S. Yang, Syuan-Ting Alex Kuo, Lauren R. Blankenship, Yan J. Sheng, Banumathi Sankaran, Pingwei Li, Carol A. Fierke, David H. Russell, Xin Yan, Shiqing Xu, Wenshe Ray Liu

## Abstract

As the COVID-19 pathogen, SARS-CoV-2 relies on its main protease (M^Pro^) for pathogenesis and replication. During the crystallographic analyses of M^Pro^ crystals that were exposed to the air, a uniquely Y-shaped, S-O-N-O-S-bridged posttranslational crosslink that connects three residues C22, C44, and K61 at their side chains was frequently observed. As a novel posttranslational modification, this crosslink serves as a redox switch to regulate the catalytic activity of M^Pro^, a demonstrated drug target of COVID-19. The formation of this linkage leads to a much more opened active site that can be potentially targeted for the development of novel SARS-CoV-2 antivirals. The inactivation of M^Pro^ by this crosslink indicates that small molecules that lock M^Pro^ in the crosslinked form can be potentially used with other active site-targeting molecules such as paxlovid for synergistic effects in inhibiting the SARS-CoV-2 viral replication. Therefore, this new finding reveals a unique aspect of the SARS-CoV-2 pathogenesis and is potentially paradigm-shifting in our current understanding of the function of M^Pro^ and the development of its inhibitors as COVID-19 antivirals.

---

SARS-CoV-2 is the viral pathogen of COVID-19 that has ravaged the whole world for more than two years. Effective vaccines that target the membrane Spike protein of SARS-CoV-2 have been developed and widely adopted for human immunization.^1^ However, the continuous emergence of new SARS-CoV-2 variants with mutations at Spike has led to viral evasion of vaccines and consequently infection surges.^2^ The situation has called for the search for SARS-CoV-2 antivirals and led to the recent success of the development of Pfizer’s paxlovid.^3,4^ SARS-CoV-2 is an RNA virus with a positive-sense RNA genome. It contains a big 20-kb open reading frame ORF1ab that is translated alternatively to form two large polypeptides pp1a and pp1ab in infected human cells.^5,6^ These two polypeptides need to be hydrolyzed to form 16 nonstructural proteins (nsps) that are key to viral biology including the formation of the RNA-dependent RNA polymerase complex for the replication of the viral genome and subgenomic RNAs. The maturation of pp1a and pp1ab is catalyzed by two internal nsp fragments. One of them is main protease (M^Pro^) that processes 13 out of the total 16 nsps.^7^ Paxlovid is a combination therapy with two chemical components. One component is nirmatrelvir that potently inhibits M^Pro^ to prevent viral replication in infected human cells.^4^

As an essential enzyme for SARS-CoV-2, M^Pro^ is an established target for the development of SARS-CoV-2 antivirals.^8-12^ Many academic research groups and a number of pharmaceutical companies have been working on the development of M^Pro^ inhibitors.^13-24^ A large international COVID Moonshot project for the development of M^Pro^ inhibitors has also been organized.^25^ As a key tool for the structure-based drug discovery, X-ray protein crystallography has been frequently employed to determine structures of M^Pro^-inhibitor complexes to assist further inhibitor optimization. This is also the approach that we have adopted for the development of M^Pro^ inhibitors.^26^ To facilitate a quick determination of M^Pro^-inhibitor complexes, we have been crystalizing the apo form of M^Pro^ and then soaking the crystals with different inhibitors for the ensuing X-ray protein crystallographic analysis.^26-28^ Our obtained apo-M^Pro^ crystals have a C121 or I121 space group that contains an M^Pro^ dimer or monomer, respectively, in an asymmetric unit. In the I121 space group, two monomers from two asymmetric units form a tightly bound dimer. In both C121 and I121 space groups, a tightly bound dimer interacts with other M^Pro^ dimers within the crystals at two regions, aa53-60 and aa216-222. The aa216-222 region is located in the noncatalytic *C*-terminal domain. Although the aa53-60 region is within the catalytic *N*-terminal domain, it is distant to the active site (Figure 1A).^26^ The M^Pro^ dimer in these crystals has an open active site whose structure rearrangements are not expected to be limited by the protein packaging in crystals (Supplementary Figure 1). Therefore, it is optimal for a soaking-based structural determination of M^Pro^-inhibitor complexes. This relatively open form of M^Pro^ is considered more desired than a closed form in representing M^Pro^ in solutions as well since M^Pro^ dimers in solutions are not expected to interact with each other. Therefore, we have been using these apo-M^Pro^ crystals and soaking them with different inhibitors for the determination of crystal structures of more than 30 M^Pro^-inhibitor complexes. One unique observation that we have made for almost all of our determined M^Pro^-inhibitor structures was a poorly defined conformation for the aa46-50 region. This region is part of the M^Pro^ active site, serving as a cap of the S2 site to bind the P2 residue in a peptide substrate (Figure 1A).^26,29,30^ The electron density of this region was very weak in all determined M^Pro^-inhibitor structures indicating a highly flexible conformation (Figure 1B and Supplementary Figure 2). There have been a number of M^Pro^-inhibitor complexes whose crystal structures have been determined and deposited into the Protein Data Bank.^8-24^ Many have a clearly defined conformation for aa46-50. However, crystals for most of these structures either had a closed form around the active site due to the protein packaging in crystals or were obtained by co-crystallization with ligands. We reason that in the closed form of the M^Pro^ crystals the protein packaging limits the structure rearrangement of aa46-50 and prevents it from adopting a flexible conformation. However, in solutions this limitation does not apply. This structural rearrangement is apparently related to the soaking process since the apo-M^Pro^ crystals that did not undergo soaking show a well-defined confirmation for aa46-50 and M^Pro^-inhibitor complexes in co-crystalized crystals did not show this rearrangement either.^8-10,26^

**Figure 1.**
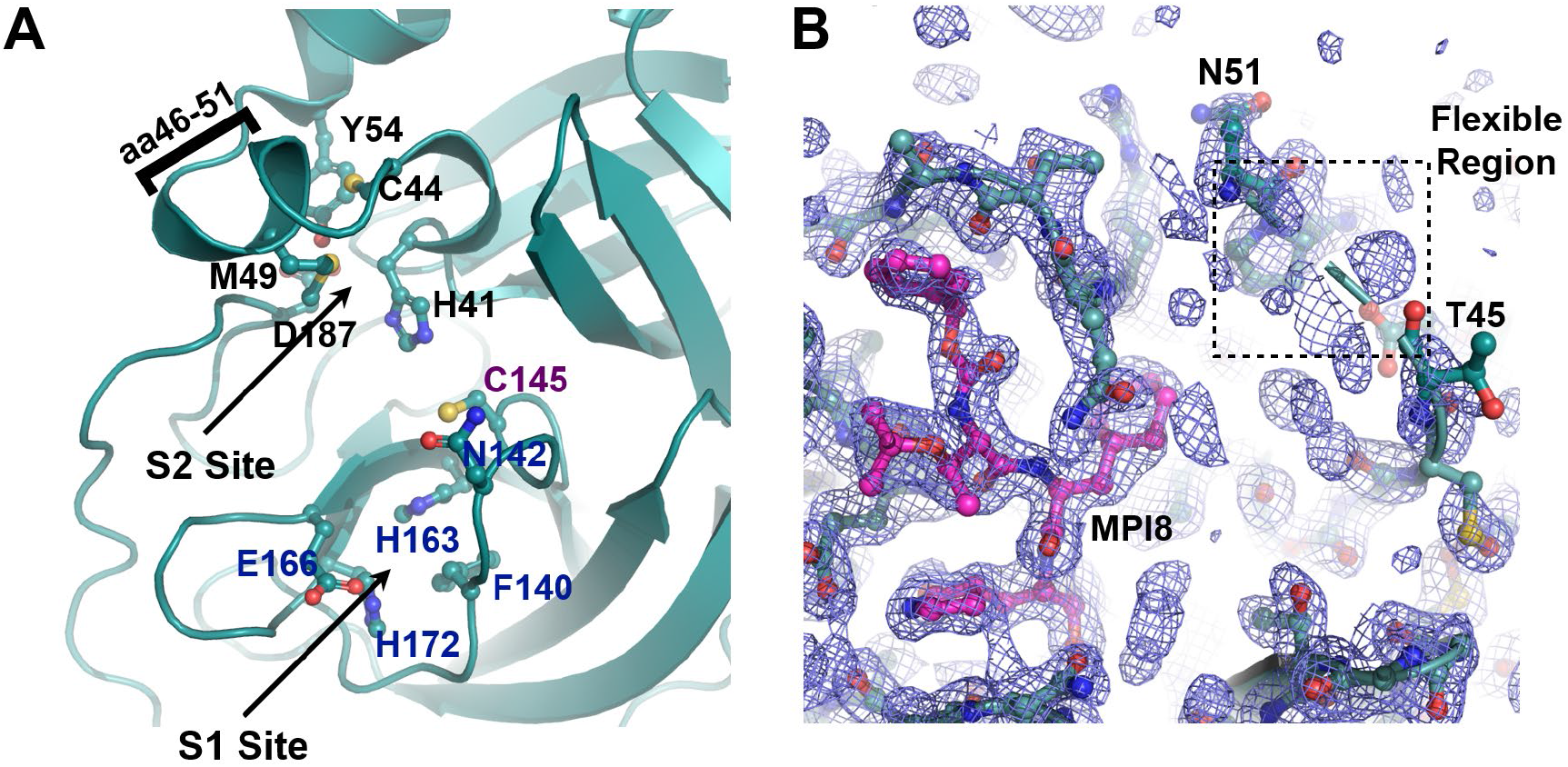
(**A**) The M^Pro^ active site based on the PDB entry 7JPY. C145 is the catalytic cysteine. Residues that form the binding sites for P1 and P2 residues in a substrate are labeled. (**B**) The low electron density map around aa46-50 in a representative M^Pro^-inhibitor complex. The ligand MPI8 that is covalently bound to C145 is shown in a color code in which carbon atoms are shown in red. For the protein, carbon atoms are shown in light teal. The aa46-50 flexible region is indicated in a dashed square. The electron density was contoured at 1*σ*.

**Figure 2.**
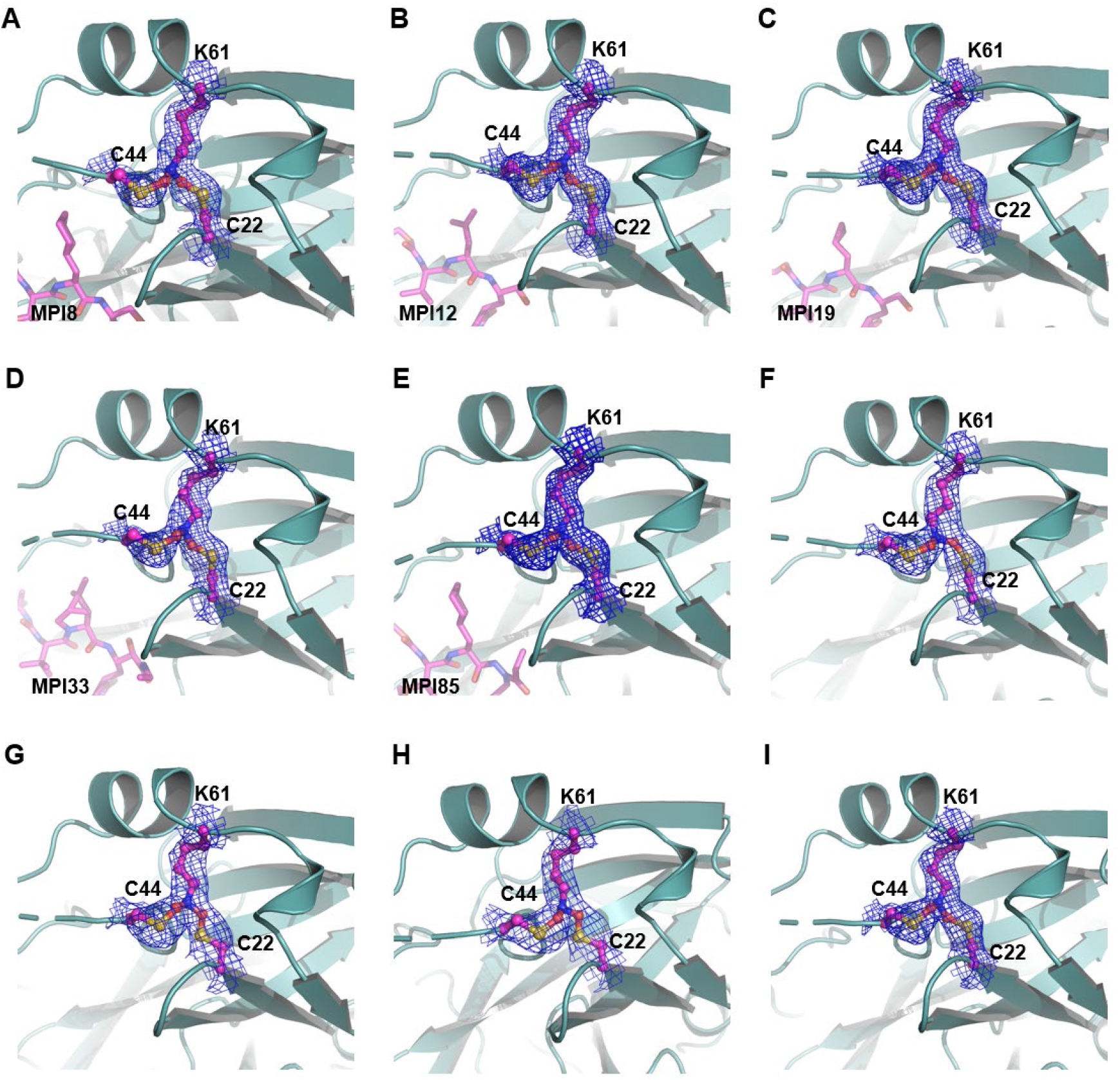
Five M^Pro^-ligand complexes and four apo enzyme structures show a SONOS-bridged crosslink between three residues including C22, C44 and K61. The electron density was contoured at 1 *σ*. The structures information is provided in Table S1.

M^Pro^-MPI8 was an early determined M^Pro^-inhibitor complex structure.^26^ In this crystal structure, besides a structurally undefined aa46-50 region, we noticed the uniquely connected electron density at residues C22, C44 and K61. As shown in Figure 2A, 2Fo-Fc electron density contoured at 1*σ*around the ends of side chains from all three residues clearly merged indicating that the three residues were covalently connected to form a Y-shaped tri-residue crosslink. Directly connecting three residues at their terminal heteroatoms to form a Y-shaped S-N-S crosslink did not fit the calculated electron density. Although there was no literature report about a Y-shaped crosslink between one lysine and two cysteine residues in a protein, a recent *Nature* article by Wensien *et al*. described a novel posttranslational N-O-S-bridged lysine-cysteine crosslink as a redox switch for the transaldolase enzyme from *Neisseria gonorrhoeae*.^31^ *N. gonorrhoeae* transaldolase undergoes oxidation to form an N-O-S crosslink in which an oxygen atom covalently bridges the K8 side chain amine and C38 side chain thiol. The consequential oxidized enzyme has a substantially lower activity than the reduced form. As a newly identified posttranslational modification, the N-O-S crosslink via oxidation serves as a regulatory mechanism for *N. gonorrhoeae* transaldolase. Since in M^Pro^-MPI8, the K61 side chain amine connected with two side chain thiol groups from C22 and C44, respectively, to form a Y-shaped crosslink instead of a linear S-O-N crosslink as observe between K8 and C38 in *N. gonorrhoeae* transaldolase, we suspected that a S-O-N-O-S-bridged, Y-shaped tri-residue crosslink in which two oxygen atoms bridged two cysteine thiol groups separately with the K61 side chain amine existed in our M^Pro^-MPI8 structure. Adding this crosslink into the structure for undergoing the structure refinement fit the calculated electron density perfectly.

Among about 10 early crystal structures of M^Pro^-inhibitor complexes, M^Pro^-MPI8 was the only one that contained the Y-shaped crosslink. However, this crosslink was subsequently also observed in several M^Pro^-inhibitor complexes determined later. Figures 2B-E show crystal structures for four other M^Pro^ complexes with MPI12, MPI19, MPI33, and MPI85, respectively. They all show a well-defined Y-shaped, S-O-N-O-S-bridged crosslink between K61 and two cysteine residues C22 and C44. We suspected that the inhibitor binding process might have triggered the formation of the crosslink, however, a careful search of our other determined structures revealed four apo-M^Pro^ structures that contained this crosslink as well. As shown in Figures 1F-H, all four structures showed clearly connected electron density at the ends of three residues C22, C44 and K61. These four apo-M^Pro^ structures were determined from apo-M^Pro^ crystals that were soaked with chemical fragments but no bound ligands were observed. So far, all structures that showed this crosslink were determined from crystals that were exposed to the air due to the soaking process.

We also determined X-ray crystal structures for several apo-M^Pro^ crystals that were not exposed to the air. They all had a structurally defined aa46-50 region and exhibited no crosslink between C22, C44 and K61. One of these structures was previously deposited into the Protein Data Bank (PDB entry: 7JPY).^26^ As shown in Figure 3A, the superposition of 7JPY over an apo-M^Pro^ structure with the Y-shaped crosslink reveals a large structural rearrangement at the region aa43-52 between two structures. In 7JPY, the side chain of C44 points toward the active site. The small aa46-51 helix tucks C44 toward the active site and makes it a key component to form the S2 binding pocket for a substrate P2 residue. The thiol group of C44 is 9.1Å away from the K61 side chain amine (Figure 3A in lemon). This distance is too far to form any possible direct interaction between the two residues. In order for C44 to physically meet K61 for a covalent interaction, the C44 backbone α-carbon moved 4.1Å closer to K61 and rotated its side chain almost 180° to adopt a conformation to form a covalent adduct with the K61 side chain amine as shown in the crosslinked apo-M^Pro^ (Figure 3A in light teal). This structural rearrangement also pushed the whole aa43-52 region away from the M^Pro^ active site, making the active site much more open. In order for C44 to translocate and rotate, a large structural rearrangement for the whole aa43-52 region is required. This explains why the Y-shaped S-O-N-O-S crosslink was observed in our crystal structures but not in most other determined M^Pro^ structures since the protein packaging in our M^Pro^ crystals allowed the large structure rearrangement at aa43-52. In 7JPY, C22 and K61 are in a close distance to each other. This close distance will likely make them form an N-O-S-bridged crosslink first and then a structurally flipped C44 will be engaged to generate the second N-O-S bridge. Since all our M^Pro^ structures that contained the Y-shaped posttranslational crosslink were determined from crystals that were exposed to the air, the molecular oxygen was most likely the reagent that generated the Y-shaped S-O-N-O-S bridge. Although how exactly the molecular oxygen oxidizes the three residues to form the crosslink needs to be further explored and confirmed, we propose a likely mechanism as shown in Figure 3B. In this mechanism, the C22 thiolate reacts with oxygen to form a peroxysulfane that then reacts with lysine to generate the first S-O-N crosslink. After C44 flips toward K61, its thiolate can react with oxygen to form the second peroxysulfane that then reacts with the first S-O-N crosslink to form the Y-shaped, S-O-N-O-S-bridged tri-residue crosslink.

**Figure 3.**
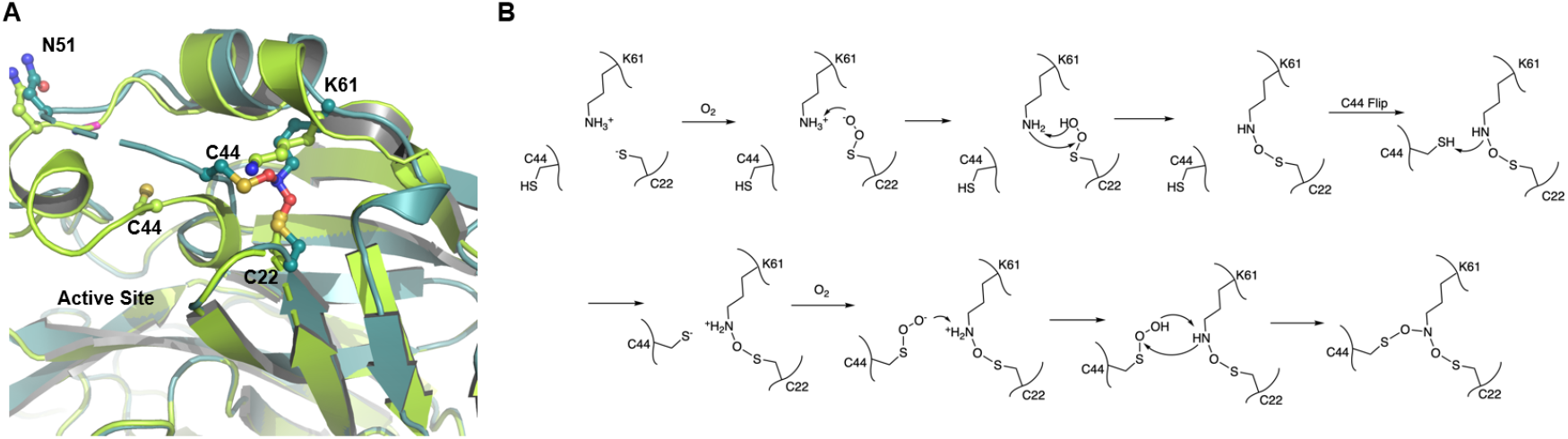
(**A**) The superposition of two different apo-M^Pro^ structures. One doesn’t have the S-O-N-O-S-bridged triresidue crosslink (lemon) and the other one does (light teal). (**B**) A proposed scheme for the formation of the SONOS-bridged triresidue crosslink in M^Pro^.

Since the crosslink formation generates a much more opened active site in M^Pro^, it presumably weakens the binding of a peptide substrate to M^Pro^ and consequently leads to a less active enzyme. To test this prospect, we freshly expressed M^Pro^ and purified it without adding a reducing reagent in lysis and purification buffers. 1 mM DTT was previously used to maintain *N. gonorrhoeae* transaldolase in its reduced state presumably due to its reduction of the N-O-S bond.^31^ We added 1 mM DTT to the freshly purified M^Pro^ as well for 30 min to reduce its potentially generated Y-shaped, S-O-N-O-S-bridged tri-substrate crosslink. We then tested the activities of the enzyme in two conditions, one with the addition of 1 mM DTT and one without. The results, as shown in Figure 4A, clearly indicate that the addition of DTT leads to about a 60% increase in activity in M^Pro^ indicating that the S-O-N-O-S crosslink is potentially a redox switch. Since there are no information such as chemical stability about the S-O-N-O-S crosslink in the literature, we conducted a native mass spectrometry analysis of the freshly purified M^Pro^ to keep the potential S-O-N-O-S crosslink intact. Our collected spectrum of M^Pro^ at 25ºC clearly showed the M^Pro^ dimer, the M^Pro^ dimer with two additional oxygen atoms, and the M^Pro^ dimer with four additional oxygen atoms (Figure 4B). The M^Pro^ dimer with four additional oxygen atoms is equivalent to M^Pro^ with the Y-shaped crosslink. The M^Pro^ dimer with two additional oxygen atoms is likely an intermediate state that has either just one M^Pro^ monomer with the S-O-N-O-S crosslink or both M^Pro^ monomers with just the S-O-N crosslink. An interesting observation was also made during the native mass spectrometry analysis of M^Pro^ when we did the in-instrument heating of M^Pro^ by sequentially increasing the temperature by 5ºC and waiting for 30 min before the native mass spectrometry analysis was conducted. When the temperature was increased to 40ºC, the majority of the M^Pro^ dimer was converted to the one containing four additional oxygen atoms. One likely explanation is that the protein at 40ºC has enhanced structural dynamics that assists the formation of the Y-shaped S-O-N-O-S crosslink. Therefore, both our activity and native mass spectrometry results support that the Y-shaped, S-O-N-O-S-bridged tri-residue crosslink potentially serves as a redox switch to regulate the SARS-CoV-2 M^Pro^ activity.

**Figure 4.**
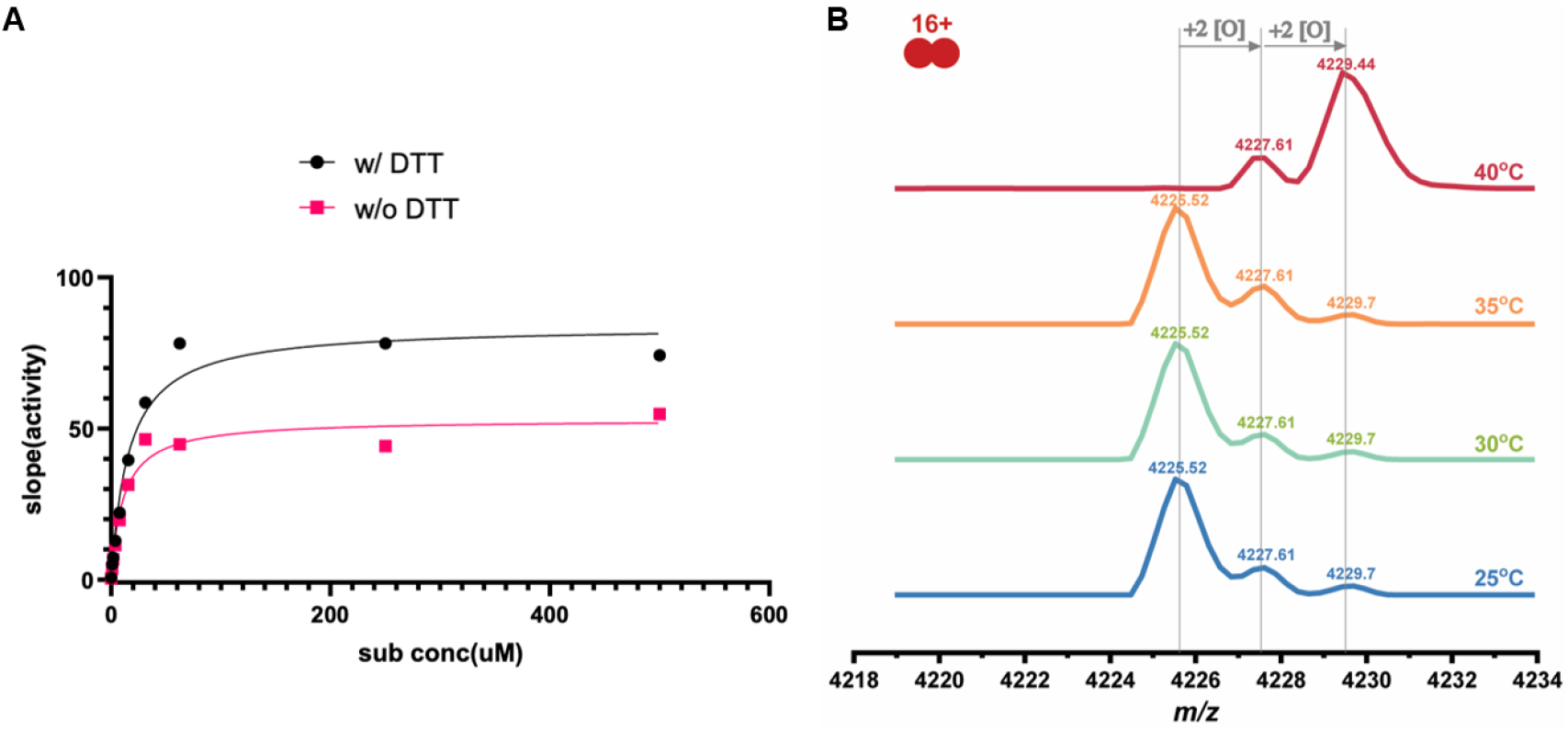
(**A**) The activity of M^Pro^ that was freshly purified without a reducing reagent in lysis and purification buffers measured in the presence (K_m_: 11 ± 2 *u*M) and absence (K_m_: 16 ± 3 *u*M) of 1 mM DTT. (**B**) Native mass spectra of M^Pro^ that was freshly purified without a reducing reagent in lysis and purification buffers and sequentially treated in the instrument by increasing the temperature by 5ºC and waiting for 30 min.

In summary, we have observed a uniquely Y-shaped, S-O-N-O-S-bridged crosslink between three residues C22, C44 and K61 in the SARS-CoV-2 M^Pro^. As a redox switch and novel posttranslational modification, this crosslink regulates the SARS-CoV-2 M^Pro^. The discovery of this crosslink is significant in that the S-O-N-O-S crosslink is a novel posttranslational modification and this novel crosslink potentially provides a key information in understanding the SARS-CoV-2 pathogenesis. Many COVID-19 patients have experienced damage to their lungs that induces hypoxia.^32^ A hypoxic condition that provides a reducing environment will favor more active M^Pro^ that drives the SARS-CoV-2 reproduction in infected human lungs. Our discovery may partially explain the severe lung damage in COVID-19 patients since the more active M^Pro^ enhances SARS-CoV-2 reproduction and therefore aggravates the symptom. SARS-CoV-2 patients will typically develop inflammation and redox imbalance with altered levels of metabolites and antioxidants.^33^ There have been many proposals to use redox-based therapeutics to potentially treat COVID-19 patients. One typical suggestion is to take antioxidants. However, the intake of antioxidants could further activate M^Pro^.^34^ Our current discovery calls into question whether taking antioxidants will be definitely beneficial to COVID-19 patients. Our finding is also important in assisting the development of M^Pro^ inhibitors as SARS-CoV-2 antivirals. The generation of the crosslink leaves a more open, larger active site allowing for the design and testing of many new inhibitors. One significant benefit of inhibitors that target this more open active site is their synergy in driving and stabilizing the crosslink formation. Since the crosslink leads to a less active enzyme, small molecules that could permanently lock C22, C44 and K61 in their crosslinked form in M^Pro^ will lead to the demise of SARS-CoV-2. Exploring this direction will allow the development of a totally new group of SARS-CoV-2 antivirals that can be provided to COVID-19 patients together with active site-targeting M^Pro^ inhibitors, including paxlovid, for potential synergistic effects in killing the virus.

## ACKNOWLEDGMENT

The work was supported by Welch Foundation (Grants A-1715 and A-2089), National Institutes of Health (Grants R21AI164088, R35GM143047, P41GM128577, and R01GM138863), and the Texas A&M X Grants Mechanism. The ALS-ENABLE beam-lines are supported in part by the National Institutes of Health, National Institute of General Medical Sciences, grant P30 GM124169-01 and the Howard Hughes Medical Institute. The Advanced Light Source is a Department of Energy Office of Science User Facility under Contract No. DE-AC02-05CH11231.

## Supplemental Information

**Figure S1.**
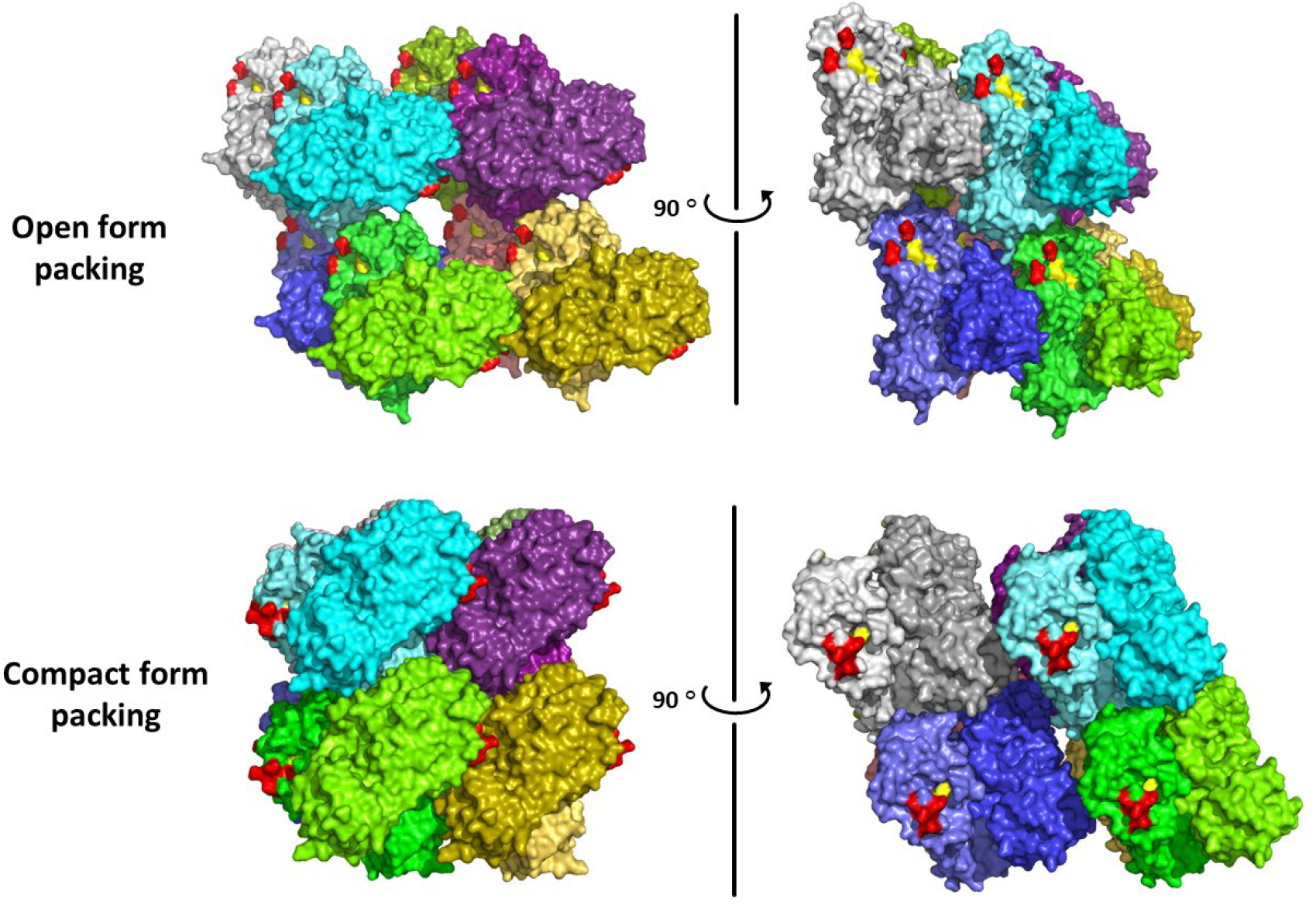
Eight dimers of M^Pro^ are shown to present the packing forms in crystals. Similar colors are given to the homogeneous monomer in one dimer. The catalytic dyad C145 and H41 are labeled as yellow. Region of aa46-50 is labeled as red.

**Figure S2.**
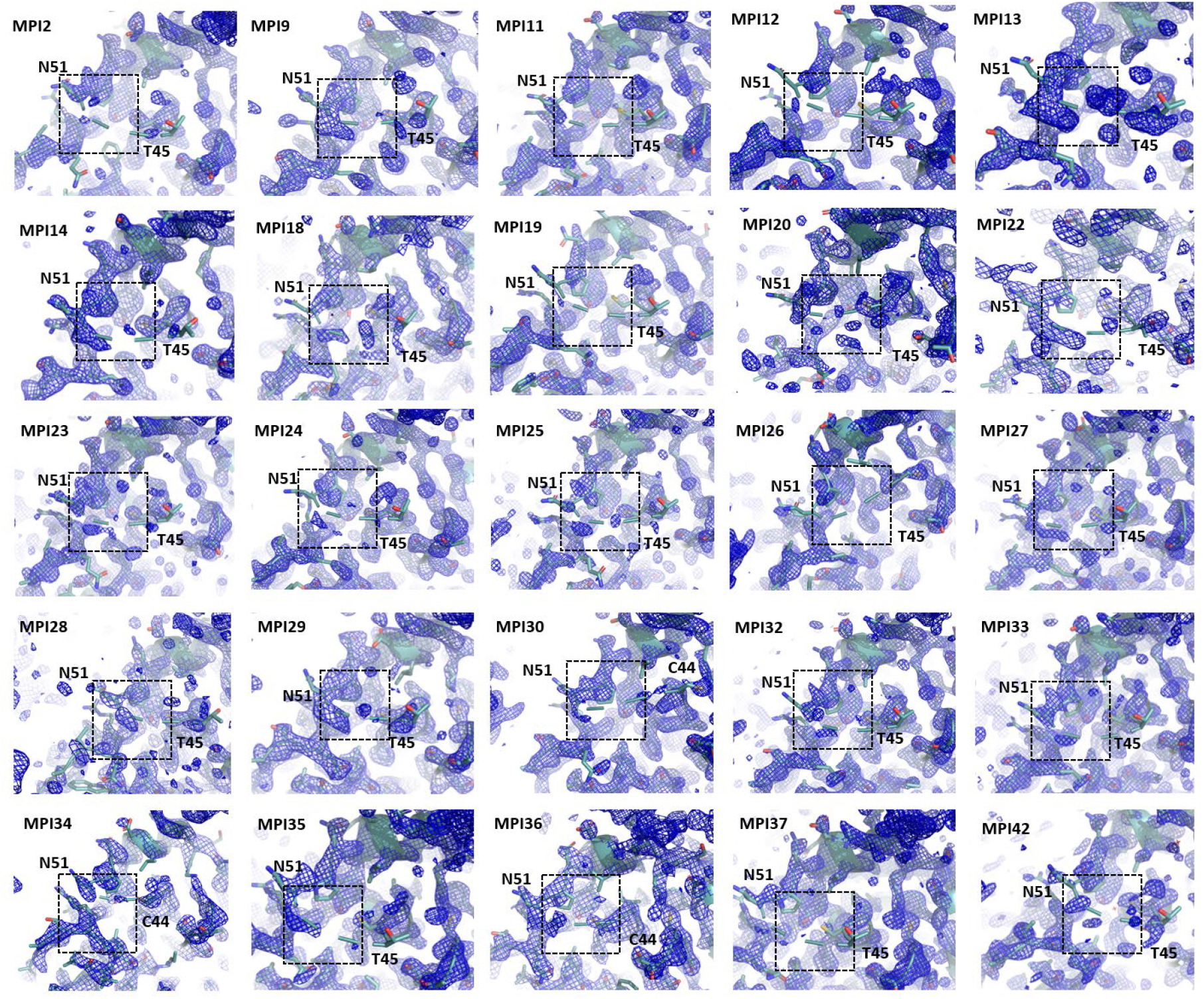
The low electron density map around aa46-50 in a representative M^Pro^-inhibitor complexes.

**Figure S3.**
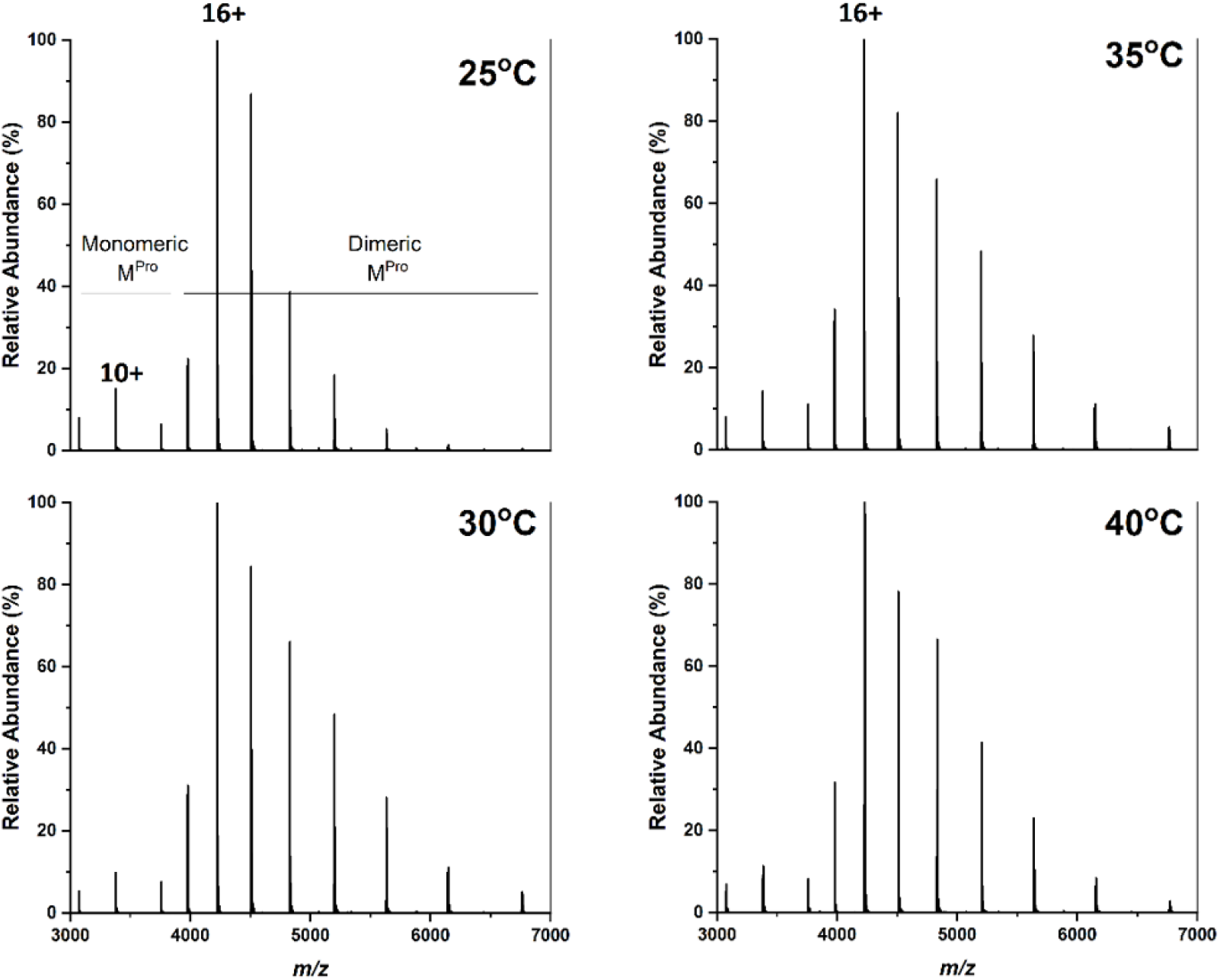
Full mass spectra of M^Pro^ at T = 25, 30, 35, and 40°C. All the spectra were collected with the same borosilicate glass tip to reduce the possible variability from tips to tips. The charge states were calculated on the basis of theoretical mass of M^Pro^ (Molecular weight 33,792) and the resolved charge state envelop. Both monomeric (with 10+ as most abundant species) and dimeric (with 16+ as most abundant species) M^Pro^ existed given the concentration of 4 μM. The 2[O] mass difference shown in the main text was calculated by using M^Pro^ dimeric form with 16+ charges.

**Table S1.** Statistics of crystallographic analysis of M^Pro^ in apo form and in complexed with inhibitors.

| Protein/Ligand<br>(PDB entry) | apo1<br>(7UU6) | apo2<br>(7UU7) | apo3<br>(7UU8) | apo4<br>(7UU9) | MPI8<br>(7UUA) |
| --- | --- | --- | --- | --- | --- |
| Data Collection |  |  |  |  |  |
| Space group | I 1 2 1 | I 1 2 1 | I 1 2 1 | I 1 2 1 | I 1 2 1 |
| cell dimensions |  |  |  |  |  |
| *a*, *b*, *c* (Å) | 51.6293 82.4004 88.6871 | 51.9582 83.3449 90.1396 | 52.1505 83.6505 89.3834 | 51.5037 83.059 87.8969 | 54.2471 80.8179 85.7217 |
| $\alpha$, $\beta$, $\gamma$ (°) | 90 96.7536 90 | 90 96.8018 90 | 90 95.9493 90 | 90 96.1827 90 | 90 96.9239 90 |
| Resolution (Å) | 24.3 - 1.85 (1.916 - 1.85) | 24.6 - 2.491 (2.58 - 2.491) | 24.56 - 2.5 (2.589 - 2.5) | 24.22 - 2.474 (2.563 - 2.474) | 24.09 - 1.85 (1.916 - 1.85) |
| *R*<sub>merge</sub> | 0.117 (1.589) | 0.078 (0.255) | 0.118 (0.631) | 0.112 (0.631) | 0.100 (1.261) |
| *I*/*σ**I* | 14.1 (0.6) | 19.4 (4.7) | 13.3 (2.4) | 14.2 (2.5) | 13.8 (1.6) |
| Completeness (%) | 95.30 (92.08) | 97.72 (82.61) | 99.32 (94.84) | 99.01 (92.42) | 99.39 (99.23) |
| Redundancy | 17.4 (5.2) | 9.5 (6.1) | 9.4 (6.6) | 9.7 (7.1) | 11.3 (8.0) |
| Refinement |  |  |  |  |  |
| No. Reflections | 30060 (2917) | 13126 (1107) | 13211 (1267) | 13089 (1219) | 31202 (3106) |
| *R*<sub>work</sub>/*R*<sub>free</sub> | 0.2239 / 0.2526 | 0.1944 / 0.2466 | 0.2021 / 0.2584 | 0.2012 / 0.2500 | 0.2435 / 0.2731 |
| No. atoms |  |  |  |  |  |
| Protein | 2360 | 2360 | 2360 | 2360 | 2327 |
| Water | 154 | 81 | 67 | 68 | 234 |
| *B* factors |  |  |  |  |  |
| Protein | 36.02 | 33.85 | 41.37 | 41.35 | 28.2 |
| Water | 39.13 | 34.19 | 39.5 | 38.99 | 32.59 |
| R.m.s deviations |  |  |  |  |  |
| Bond lengths (Å) | 0.011 | 0.009 | 0.015 | 0.01 | 0.01 |
| Bond angles (°) | 1.16 | 1.07 | 1.26 | 1.22 | 1.06 |
| Protein/Ligand<br>(PDB entry) | MPI12<br>(7UUB) | MPI19<br>(7UUC) | MPI33<br>(7UUD) | MPI85<br>(7UUE) |  |
| Data Collection |  |  |  |  |  |
| Space group | I 1 2 1 | I 1 2 1 | I 1 2 1 | I 1 2 1 |  |
| cell dimensions |  |  |  |  |  |
| *a*, *b*, *c* (Å) | 51.92 81.18 90.0856 | 52.01 81.32 89.4029 | 54.2617 81.3062 85.6688 | 54.6222 83.3566 85.8644 |  |
| $\alpha$, $\beta$, $\gamma$ (°) | 90 96.2965 90 | 90 96.1217 90 | 90 97.4294 90 | 90 96.8577 90 | |
| Resolution (Å) | 47 - 1.63 (1.688 - 1.63) | 46.93 - 1.6 (1.657 - 1.6) | 23.82 - 1.85 (1.916 - 1.85) | 24.73 - 1.85 (1.916 - 1.85) |  |
| *R*<sub>merge</sub> | 0.066 (0.776) | 0.050 (2.064) | 0.071 (0.701) | 0.084 (2.043) |  |
| *I*/*σ**I* | 12.5 (1.6) | 11.3 (0.8) | 19.4 (2.8) | 16.6 (1.3) |  |
| Completeness (%) | 95.46 (91.40) | 99.56 (99.81) | 95.77 (94.30) | 93.47 (83.22) |  |
| Redundancy | 5.9 (5.5) | 4.8 (4.8) | 11.8 (8.3) | 10.4 (7.4) |  |
| Refinement |  |  |  |  |  |
| No. Reflections | 44234 (4239) | 48554 (4840) | 30206 (2961) | 30516 (2698) |  |
| *R*<sub>work</sub>/*R*<sub>free</sub> | 0.2093 / 0.2281 | 0.2082 / 0.2292 | 0.2089 / 0.2356 | 0.2041 / 0.2314 |  |
| No. atoms |  |  |  |  |  |
| Protein | 2347 | 2347 | 2338 | 2338 |  |
| Water | 199 | 149 | 221 | 232 |  |
| *B* factors |  |  |  |  |  |
| Protein | 34.53 | 42.18 | 26.81 | 28.03 |  |
| Water | 40.9 | 48.51 | 32.2 | 34.32 |  |
| R.m.s deviations |  |  |  |  |  |
| Bond lengths (Å) | 0.01 | 0.013 | 0.01 | 0.01 |  |
| Bond angles (°) | 1.29 | 1.93 | 1.12 | 1.08 |  |

### MATERIALS AND METHODS

#### Recombinant M^Pro^ protein expression and purification

The pET28a-His-SUMO-M^Pro^ expression and purification are according to our previous report.^35^ The pET28a-His-SUMO-M^Pro^ construct was transformed into *E. coli* BL21(DE3) cells. Transformed cells were cultured at 37°C in 2xYT medium with kanamycin (50 g/mL) until OD_600_ reaching 0.6, then induced with 1 mM isopropyl-D-1-thiogalactoside (IPTG) at 37 °C. After 3 hours, cells were harvested and lysed in buffer A (20 mM Tris, 100 mM NaCl, 10 mM imidazole, pH 8.0). The supernatant was loaded onto a nickel-chelating column (GenScript) washed with buffer A, followed by elution with buffer B (20 mM Tris, 100 mM NaCl, 250 mM imidazole, pH 8.0). The eluted protein solution was desalted to buffer C (20 mM Tris, 10 mM NaCl, pH 8.0) by HiPrep 26/10 desalting column (GE Healthcare). The His-SUMO-M^Pro^ proteins were digested with SUMO protease overnight at 4°C. The digested protein was applied to nickel-chelating column again to remove the His-tagged SUMO protease, the His-SUMO tag, and the expressed protein with uncleaved His-SUMO tag. The tag-free M^Pro^ protein was loaded to the size exclusion column HiPrep 16/60 Sephacryl S-100 HR (GE Healthcare) pre-equilibrated with buffer D (20 mM Tris, 100 mM NaCl, 1 mM EDTA, pH 7.8). The eluted M^Pro^ protein was stored with buffer D in -80 °C for further use.

#### Km analysis of M^Pro^

The assays were carried out with 50 nM M^Pro^ at 37°C with continuous shaking. The Sub3 substrate (DABCYL-Lys-Thr-Ser-Ala-Val-Leu-Gln-Ser-Gly-Phe-Arg-Lys-Met-Glu-EDANS) was purchased from BACHEM and stored as 10 mM solution in 100% DMSO at -80°C. Enzyme activity was monitored by detecting fluorescence with excitation at 336 nm and emission at 490 nm wavelength. The dilution buffer (used for enzyme and substrate dilution) is 10 mM Na_x_H_y_PO_4_, 10 mM NaCl, 0.5 mM EDTA, pH 7.6. To conduct the Km analysis, the enzyme was diluted to 100 nM using dilution buffer with or without 1 mM DTT and set on ice for 30 min. 25 μL of the 100 nM enzyme solution was then added to a 96-well plate and incubated at 37°C for 30 min. When the incubation period was over, 25 μL of each substrate solution, final concentration from 4 μM to 500 μM with 5% DMSO, was added to each well to initiate the assay. The first 0-200 seconds were analyzed by linear regression for initial slope analyses with GraphPad Prism 8.0.

#### X-Ray Crystallography Analysis

The production of crystals of apo M^Pro^ and M^Pro^-inhibitor complexes was following the previous protocols with the crystal growing condition of 0.1 M Bis-tris, pH6.5, 16% w/v PEG10k.^35^ The data of M^Pro^ with MPI12 was collected on a Rigaku R-AXIS IV++ image plate detector, the data of M^Pro^ with MPI19 was collected at the Advanced Light Source (ALS) beamline 5.0.2 using a Pilatus3 6M detector, all the other data were collected on a Bruker Photon II detector. The diffraction data were indexed, integrated and scaled with iMosflm or PROTEUM3^36^. The structure was determined by molecular replacement using the structure model of the free enzyme of the SARS-CoV-2 M^pro^ [Protein Data Bank (PDB) ID code 7JPY] as the search model using Phaser in the Phenix package ^35,37^. *JLigand* and *Sketcher* from the CCP4 suite were employed for the generation of PDB and geometric restraints for the inhibitors. The inhibitors were built into the *Fo-Fc* density by using *Coot* ^38^. Refinement of all the structures was performed with Real-space Refinement in Phenix ^37^. Details of data quality and structure refinement are summarized in Table S1. All structural figures were generated with PyMOL (https://www.pymol.org).

#### Native Mass Spectrometry

Native mass spectrometry (nMS) analysis was performed on a Q Exactive UHMR Hybrid Quadruple-Orbitrap Mass Spectrometer (ThermoFisher) with m/z range was set from 1,000 to 10,000 and with resolution set to 12,500 (at m/z 400). 10 μL of the sample was loaded to a borosilicate glass capillary tip (Sutter, CA) with 1100 to 1500 V spray voltage supplied by an inserted platinum wire. Important parameters to reduce non-specific adducts include: capillary temperature 100oC, in-source trapping and activation -10 V, ion transfer high m/z, collision-induced dissociation (CID) 10 eV, and higher energy dissociation (HCD) 30 V. In variable-temperature electrospray ionization (vT-ESI) experiment, the temperature of solution was controlled as previously described.^39^ The temperature was varied from 25oC to 40oC with 5oC increments and the equilibrium time at each temperature was 5 minutes.

